# Stability, incumbency and ecological reorganization after the Permian-Triassic mass extinction

**DOI:** 10.1101/241638

**Authors:** Peter Roopnarine, Allen Weik, Kenneth Angielczyk, Ashley Dineen

## Abstract

The Permian-Triassic mass extinction (PTME) altered macroevolutionary landscapes by removing incumbent biota. Here, using terrestrial paleocommunities of the Karoo Basin spanning the PTME, we show that a pre-extinction incumbent configuration of biotic interactions made significant ecological re-organizations or macroevolutionary innovations unlikely. The post-PTME ecosystem initially was more likely to be reorganized, but incumbency was re-established by the Middle Triassic. We argue that the stability of the pre-PTME ecosystem, its subsequent loss, and replacement, resulted from the influence of community-level structure and dynamics on species evolution and survival.

**One sentence summary:** Biotic incumbency at the end of the Paleozoic, based on community functional organization, was destroyed by the Permian-Triassic mass extinction, allowing the development of novel community types.

## Introduction

The rise and fall of biotic interactions and incumbent, geologically persistent taxa, are key features of long-term macroevolutionary and macroecological patterns in the fossil record (*1*). Although the replacement of incumbents and interactions have been explained in various ways (*1–6*), mass extinctions played particularly important roles. During these transformative events, communities underwent significant compositional and ecological reorganizations, or were replaced completely by ecologically novel systems, giving mass extinctions an effect on history disproportionate to their contributions to the total number of Phanerozoic extinctions (*7*). Here we show, using a well-documented series of terrestrial paleocommunities spanning the Permian-Triassic mass extinction (PTME), and a numerical model of community stability (*8*), that incumbency before the PTME was maintained by a pre-existing community-level structure of biotic interactions. When compared to alternative evolutionary trajectories and ecological reorganizations, this structure was more advantageous to long-term species persistence and co-existence. The structure’s loss through successive waves of extinction resulted in an Early Triassic community in which reorganization would have significantly improved persistence and coexistence. By the Middle Triassic, this recovery stage was replaced by a community in which innovation and reorganization would have been disadvantageous, thereby forming the basis for renewed incumbency.

We studied seven paleocommunities from the Karoo Basin, South Africa, ranging from the late Permian (Wuchiapingian) lower *Daptocephalus* Assemblage Zone, to the Middle Triassic (Anisian) *Cynognathus* Assemblage subzone B (*31, 35, 36*) (Fig. 1; table S1) (*12*). Taxon composition between successive communities was nearly constant during the Permian, whereas turnover was dramatically greater in the Triassic. We modeled each paleocommunity as an ensemble of food webs of size S (number of species), with a structural complexity determined by the hierarchical and functional partitioning of S into G trophic guilds, and E sets of inter-guild interactions. The ensemble was organized as four nested sets of hypothetical communities of increasingly constrained structural complexity (*33*). The most inclusive set comprised random networks of size S, constrained by having the total number of interactions drawn from a mixed exponential power law distribution (*12*); the least inclusive consisted of the observed paleocommunity itself. Nested between these end members were: model communities with S partitioned into G guilds with E links, yielding food webs of structural complexity equal to, but compositionally different from the observed paleocommunity; and models containing the same functional structure of the paleocommunity, but with randomized partitioning of S among guilds. We thus examined the effects of imposing structural features on the ensemble that increasingly required random food webs of size S to be more consistent with observed paleocommunity structure.

**Fig. 1.**
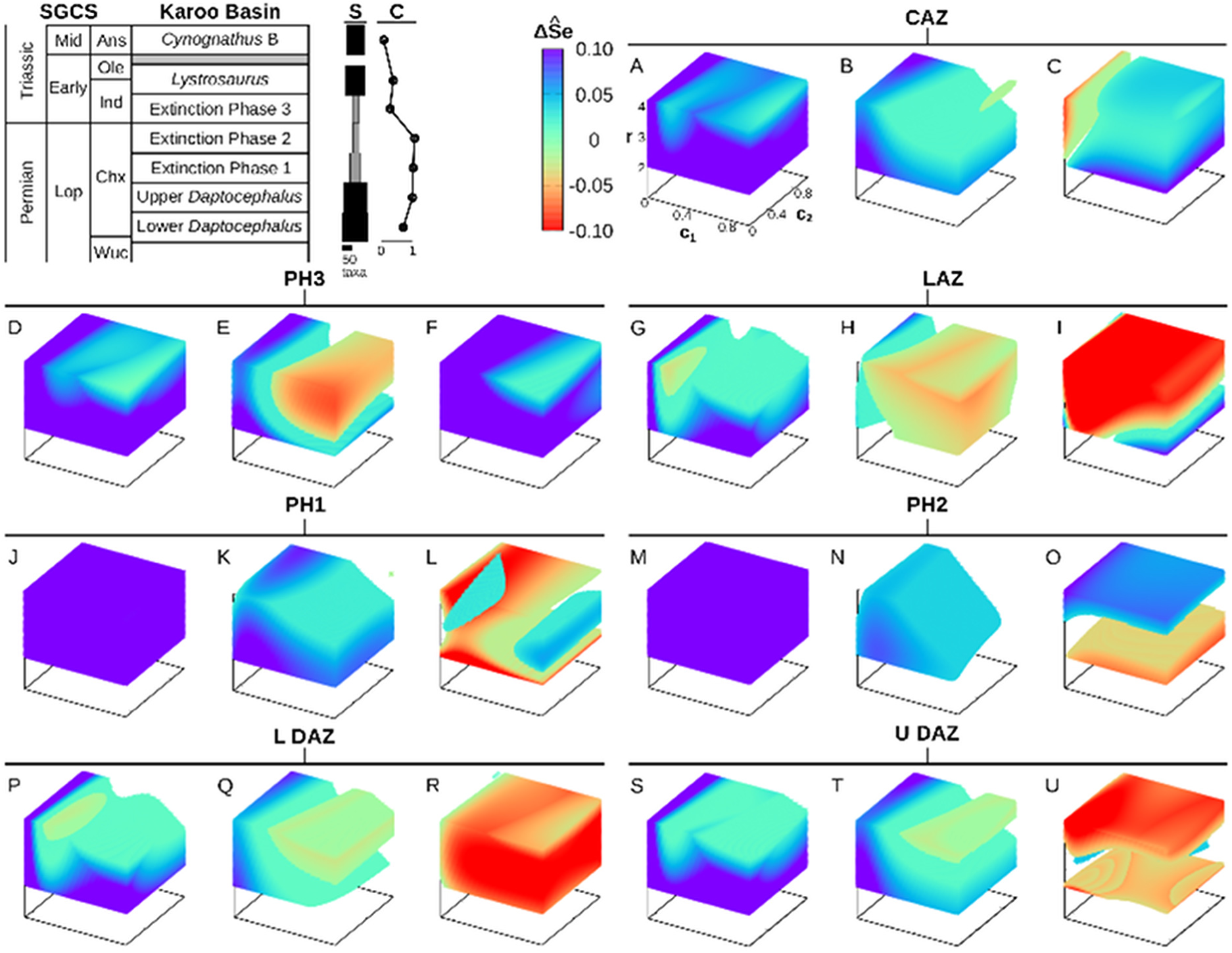
Model global stabilities and paleontological context. Upper left panel summarizes the stratigraphic position of each paleocommunity, its taxon richness (S) and taxon continuity (C). Gray bars in S represent the extinction phases. C is calculated as the fraction of vertebrate genera that have persisted from the previous community. (**A-U**) For each paleocommunity, ΔSe between the observed paleocommunity and (left) random, (middle) guild altered, and (right) richness altered manipulations. ΔSe > 0 represented as cooler colors. ΔSe = 0 uncolored. Paleocommunity acronyms: CAZ-*Cynognathus* Assemblage Zone (AZ), LAZ - *Lystrosaurus* AZ, Ph3 - Extinction Phase 3, Ph2 - Extinction Phase 2, Ph1 - Extinction Phase 1, L DAZ - lower *Daptocephalus* AZ, U DAZ - upper *Daptocephalus* AZ.

The structurally complex models represent hypothetical alternative evolutionary pathways, ranging from minimally divergent (species richnesses within guilds vary from observed because of differential rates of origination/extinction), to more divergent histories where ecological innovation, origination and extinction could result in alternative community types. If all histories were possible, incumbency would emerge if a particular type of community consistently supported a greater number of stably coexisting species, and thus greater species persistence relative to other communities. We estimated levels of coexistence, Se, associated with real and hypothetical communities using a tractable model of interspecific interactions (*8*)(*12*).

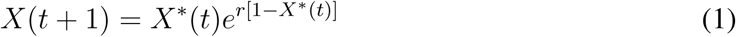

Population sizes, X, are functions of intrinsic population growth rate, r, both positive and negative interspecific trophic interactions (*β*), and normalized carrying capacity. X^*^ is population size modified by interspecific interactions as

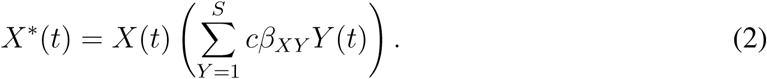

c comprise scaling factors ranging between zero and one, that modify the average interaction strengths 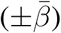 of the community. Populations become extinct because larger values of r increase the variability of population trajectories (Fig. S1, S2), and more negative interaction strengths decrease species feasibilities, but the system always settles dynamically to a stable state, yielding Se. The system thus undergoes a May-Wigner transition (*8, 14*), with Se declining nonlinearly with increasing S, number of interspecific interactions, or 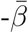(*12*) (Fig. S3).

We explored the dependence of Se on r and 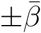 for each random, alternative or observed model of a paleocommunity by simulating Equation 1 at values of r ranging from 2-4. Average interaction strengths were drawn randomly from a uniform distribution ranging from 0-1 and were scaled by factors ranging from 0.1-1 (c in Equation 2). Thirty species level food webs were simulated for each model at each parameter set (r, −*c*_1_*β*, *c*_2_*β*), for a total of 27,000 food webs per model per community. Differences between observed and hypothetical models (ΔSe) were tested with parameter dependent t-tests (*12*), and visualized as regions of parameter space where the global stability of the models differ significantly (Fig.1A-U).

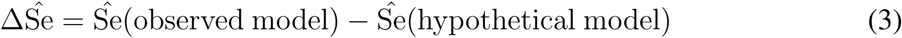

where 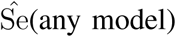 is a regression estimate of Se (Fig. S4). One community model typically did not outperform another at every parameter set (although see Fig. 1D, 1F, 1J and 1M). Specific values of intrinsic population growth rates and biotic interactions determine the circumstances under which a model may be expected to support a significantly greater number of species compared to another. By contrasting observed paleocommunity structures to random communities and alternative model communities, several important conclusions may be drawn regarding the basis for a pre-PTME pattern of incumbency, the loss of incumbency, and the ecological transformation wrought by the PTME.

First, the decline of Se with increasing population growth rates (r) or negative interaction strengths (−*β*) depends critically on the higher level, functional organization of the community. Compartmentalization into guilds slows the decline relative to random, uncompartmentalized food webs of equal S, regardless of whether compartments are observed or hypothetical. The parameter range under which compartmentalized (guild-structured) communities would support significantly more species than random communities (ΔSe > 0) depends on S and the structure of the community. During the PTME, when S was lowest, compartmentalization would have permitted greater species coexistence under all circumstances of r and *β*, relative to random communities (Fig. 1J and 1M). Although this is generally true of pre-and post-PTME paleocommunities, there are cases, when r and −*β* are both high, or −*β* has been scaled by a factor of approximately 0.2, that ΔSe ≤ 0 (Fig. 1G and 1P). Those cases represent conditions under which paleocommunity structure would have been more ecologically and evolutionarily labile; structure could be altered by changes to species properties or species composition, without a negative impact on stability.

Second, during the late Permian, observed paleocommunities would have been more globally stable than alternatively structured communities (ΔSe > 0), along a gradient of decreasing r and −*β* (Fig. 1K,1N,1Q,1T). Observed paleocommunities prior to the PTME could be less stable at greater values of r and −*β* (ΔSe < 0) (Fig.1Q,1T), whereas during the extinction, ΔSe > 0 (Fig. 1K,1N). These contrast with models that preserve guild structure, but randomize the partitioning of S among guilds (Fig. 1L, 1O, 1R, 1U). Those hypothetical communities would possess the same interacting lineages as the paleocommunities, but would be distinguished by having different diversification and extinction rates. In those cases, the hypothetical models are generally more stable than the paleocommunities themselves within much of the parameter range. Thus, the pre-existing Permian community structure was maintained by its pattern of biotically interacting lineages, but not specifically the numbers of species involved in those interactions.

Third, no patterns of incumbency could have been established in the Early Triassic. The Early Triassic third phase of the PTME comprised survivors from the Permian Karoo Basin ecosystem, and likely immigrants from neighboring regions (*15*). Structural re-organizations of this assemblage could have generated more stable and persistent alternative communities (Fig. 1E), at parameter values encompassing more realistic ecologies when compared to pre-PTME communities (Fig. 1Q, 1T). Moreover, ΔSe > 0 throughout the parameter space when compared to alternative partitionings of S (Fig. 1F), showing that although the community comprised species capable of surviving the PTME, options for subsequent within-guild variation of taxon richness were very narrow. Changes via origination and extinction within existing lineages would have yielded less stable communities, producing a macroevolutionary dead-end. The succeeding *Lystrosaurus* Assemblage Zone (LAZ) has been noted for its unusually rich amphibian fauna (*30,38*), numerical dominance of the herbivorous therapsid *Lystrosaurus* (*38*), accelerated ontogenetic development of some tetrapods (*18, 19*), and unusual ecological dynamics (*21, 32*). Our results show that both variation of guild richnesses (Fig. 1I), and reorganizations of guild structure (Fig. 1H) would have been overwhelmingly likely to result in more persistent species and stable communities than those observed. The LAZ community itself would have persisted unchanged, with lineages therefore apparently incumbent, only if negative species interactions were exceedingly weak (Fig. 1H). This contrasts with the later, Middle Triassic *Cynognathus* Assemblage Zone (CAZ), wherein any changes to guild richnesses or paleocommunity structure were unlikely to yield greater persistence or stability (Fig. 1B, 1C). The CAZ community signaled a return to incumbency in the Karoo ecosystem.

Paleocommunity compositional and structural constancy prior to the PTME resulted from greater global stability compared to communities that could have arisen via macroevolutionary variation. This incumbency was generated by the persistence of interacting lineages, even during the PTME (Fig. 1), and the communities could have accommodated changing taxon diversities while maintaining ecological functions. Successful ecological reorganization was unlikely. The extinction and replacement of most lineages led to an Early Triassic community (LAZ) that would have been more ecologically and evolutionarily variable, and labile. We posit that patterns of ecological reorganization and evolutionary innovation in the wake of mass extinctions are therefore accounted for, at least in part, by the destruction of highly stable, preexisting community structures, and the subsequent time required for the development of new systems of stably coexisting, biotically interacting lineages. The rapidity with which taxon richness increased in the wake of the PTME further suggests that recovery does not require extended intervals of time for the evolution of new biotic interactions. Instead, time is required for the evolution of persistent assemblages of interactors and interactions.

## Supplementary Materials

Materials and Methods

Supplemental Text

Table S1 -S2

Fig S1 -S4

References (22 - 47)

## Supplementary Materials for Stability, incumbency and ecological reorganization after the Permian-Triassic mass extinction

**This PDF file includes:**

Materials and Methods

Supplementary Text

Figs. S1 to S4

Tables S1 to S2

### Materials and Methods

#### Paleocommunity datasets

The rocks of the Beaufort Group of the South African Karoo Basin are highly fossiliferous (*23–25*), and provide a nearly continuous record of terrestrial community transformation from the middle Permian to the Middle Triassic (*26–29*). The dataset presented here is a modification of our previous works on the food webs of the Permo-Triassic communities of the Karoo Basin (*30–35*). We subdivided the Permian data to reflect Viglietti et al.’s (*36, 37*) subdivision of the former *Dicynodon* Assemblage Zone into upper and lower *Daptocephalus* Assemblage Zones (upper and lower DAZ). Our treatment of stratigraphic ranges, and thus community compositions, generally follow Viglietti et al. However, there are a few rare taxa that were not included in their stratigraphic range chart, making it uncertain whether they were members of the upper or lower DAZ communities. We included those taxa in both the upper and lower DAZ, recognizing that this might not reflect the true stratigraphic ranges. All data of tetrapod taxa and ranges used in this paper are available in (*32*).

Smith and Botha-Brink (*25*) recognized three phases of extinction near the Permian-Triassic boundary in the Karoo Basin. The last of these communities (Extinction Phase 3, Ph3) comprises the 10 tetrapod taxa that Smith and Botha-Brink included in their Extinction phase 3 and Recovery phase, as well as a small number of additional tetrapods known from equivalent strata at other localities not considered in Smith and Botha-Brink. The successive LAZ community does not include the five taxa that Smith and Botha-Brink indicated went extinct in the earliest Triassic (*Promoschorhynchus*, *L. curvatus*, *Tigrisuchus*, *Proterosuchus*, *Progalesaurus*), but it does include all other taxa reported from the LAZ, including those that originated after the stratigraphic range chart in Smith and Botha-Brink’s Figure 12.

Hancox et al. (*38*) proposed an informal three-fold subdivision of the Middle Triassic *Cynognathus* zone, and this subdivision has been widely adopted by subsequent workers. The *Cynognathus* B subzone is the best sampled (*39*), and represents the “classic” *Cynognathus* zone assemblage that has been recognized for over a century (*38, 40–44*). Because part of our interest is to document changes in community composition and food web structure over time in the aftermath of the PTME, we chose to focus on the *Cynognathus* B subzone in this paper, as it represents the best documented discrete community in the Karoo Basin from the time period represented by the *Cynognathus* zone.

#### Food web reconstruction

The general procedure for compiling the data underlying our trophic networks, or food web of interactions between species, and the construction of the networks themselves, are described in detail by Roopnarine et al. (*31, 32*) and Roopnarine and Angielczyk (*34, 35*), and have not been altered for the present paper. We recognized a total of 26 guilds in our paleocommunities (table S1). A guild is herein defined as a group of species utilizing the same resources in similar ways (*45*) and sharing other properties such as habitat, body-size ranges, and predators, e.g., “large amniote carnivores”, or “very small carnivores/insectivores. Species richness of each guild varies among the communities, and guilds may be vacant if the appropriate species are missing from the community. Our set of guilds includes four primary producer guilds, and both terrestrial and aquatic invertebrates and vertebrates. Most of the records in the dataset represent genera, reflecting the fact that many of the vertebrate genera are effectively monotypic. We recognized multiple species within genera where they were supported by strong taxonomic evidence and had distinct stratigraphic ranges. In most cases, tetrapod body size categories were based on skull length as follows: very small, 0-100 mm; small, 101-200 mm; medium, 201-300 mm; large, 301-400 mm; very large, 401 mm and above. Tetrapod size classes overlap strongly in the guilds on which they prey if those size classes are contiguous. The range of prey size classes is based on the fact that predators can often prey on species whose maximum body sizes exceed the maximum for that predator, but whose ontogenetic and adult size variance fall within the range of expected prey (*32*).

An important difference between the food web models constructed here and those used in previous studies lies in the treatment of plant guilds. Here, plant guilds each have a richness of one. This is necessary because the current state of paleobotanical systematics in the Karoo paleoecosystem does not allow accurate estimates of guild richness, as we have for animals. Therefore, we assume plant species within a guild to be effectively neutral (*46*), treating them as a single population, and hence producer guild richnesses of one.

##### Supplementary Text

###### In-degree distribution

Each food web within an ensemble comprised species where the number of prey species per consumer was drawn from a mixed exponential-power law distribution, P(k) (*47*), which compensates statistically for trophic interactions that are lost during the fossilization of a food web (*48*).

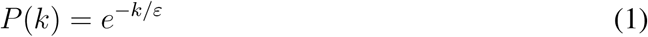

where
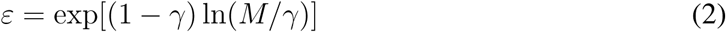

*γ* is a power law exponent (2.5 for all simulations conducted here) and M is the total species richness of all guilds that are prey to the one under consideration.

###### Population model

The exponential map is used here as a basic discrete model of population growth. A discrete model was selected to maintain tractability as the number of populations or species, S, in a system grows. For each species, its population grows as a function of an intrinsic growth rate, r, and carrying capacity, which is normalized to one in the model. Thus,
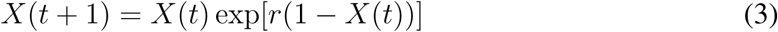

In a single species model, r determines the population dynamics, with the complexity of the dynamic increasing as r increases. The dynamic undergoes a series of transitions typical of other discrete population growth models, such as the discrete logistic map, transitioning from an initial stable point equilibrium (r ≈ 2), to stable oscillations, quasi-periodicity, and finally chaos (Supplemental Fig.S1A). The transition to chaos occurs at r ≈ 2.7 for the exponential map (Supplemental Fig.S1B).

Species predator-prey interactions are incorporated into the model by modifying the population size of each species at each step to reflect interspecific interactions.

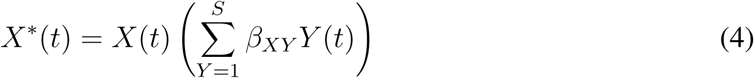

where *β* is the per capita strength of interaction between species X and Y, and will be negative if Y preys on X, and positive if X preys on Y. Equation 4 is then substituted into Equation 3 to yield a population dynamic reflective of interspecific interactions,
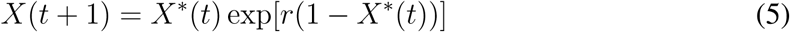

###### Food web simulations

Species-level food webs were generated from the guild level data, and P(k), as described in detail in earlier works (Roopnarine et al.,2007; Roopnarine, 2009, 2010; Roopnarine and Angielczyk, 2011, 2015). Each food web was represented as a SxS matrix of interspecific interactions, where predator-prey interactions were generated as described in the main text. Self-regulatory interactions (diagonal matrix elements) equaled zero for all species in the model. Off-diagonal elements were then modified by parameters c1 and c2 for negative and positive elements, respectively, representing the scaling of interaction strengths. The assignment of a value for r then allows Se of the web to be determined by iterating Equations 4 and 5. For each paleocommunity, random and alternative models, 30 food webs were generated at a (r,-c1*β*,c2*β*) parameter set. Overall, r ranged from 2 to 4 in increments of 0.25, and c1 and c2 ranged from 0.1 to 1 in increments of 0.1. Therefore, a total of 27,000 food webs were generated for each model. Each parameterized web was iterated for 10,000 steps, and Se measured as the number of species for which X(t) > 0.

Population trajectories within any of the food web models are representative of an array of empirically and theoretically expected forms, including stable point equilibria, stable limit cycles, quasi-periodic cycles, and chaos (Fig. S2). Quasi-periodic and chaotic trajectories were confirmed by computation of Lyapunov exponents. The transition to complex dynamics, including chaos, generally occurs at values of r as low as 2.0, which is below the expected value of 2.7 for a single species system. This highlights the fact that the major source of complex dynamics in the food webs are not species-specific characteristics; instead they are generated by asynchronous interspecific impacts, i.e. the time scales on which species affect each other are spectra of the lengths of the multiple paths that connect them.

###### Se and ΔSe

Se was measured as the number of species extant after 10,000 time steps, and for all models declines with increasing r and average negative interaction strengths (Fig. S3). The latter value is controlled by the scaling factor c1. The differences between two models, ΔSe, were tested separately at each particular parameter coordinate using a t-test comparison of the 30 simulations, of each model, at that parameter coordinate. The resulting t statistics were used to establish the range of ΔSe within which the models did not differ significantly (with significance *α* = 0.01).

The parameter dependence of Se was modeled using a nonlinear power function of the form
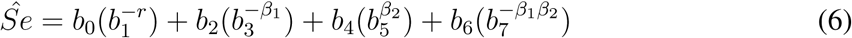

where *β*_1_ and *β*_2_ are negative and positive community matrix elements respectively. Coefficients b0-b7 were fit using ordinary least squares multiple regression. The dependence of Se on positive *β* was frequently positive, or zero (non-significant). Note the necessary inclusion of an *β*_1_*β*_2_ interaction term.

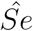 was used to create a more detailed model of the dependence of Se on parameters than was computed originally. Computational limitations arise when the resolutions of the parameter increments over which Se is calculated are increased. For example, r was evaluated over the range 2-4, in increments of 0.25. The statistical significance of Equation 6 (Table S2), however, allows us to model the behavior of Se over finer scales. 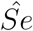 was therefore computed over the same parameter ranges, but with r, c1 and c2 being increased in increments of 0.05. This higher resolution modeling allows both a useful visualization of the behavior of Se throughout the parameter space, and a simple calculation and visualization of the difference between two models, i.e. significantly different ranges of ΔSe (Fig. S4).

---

**Fig. 1.**
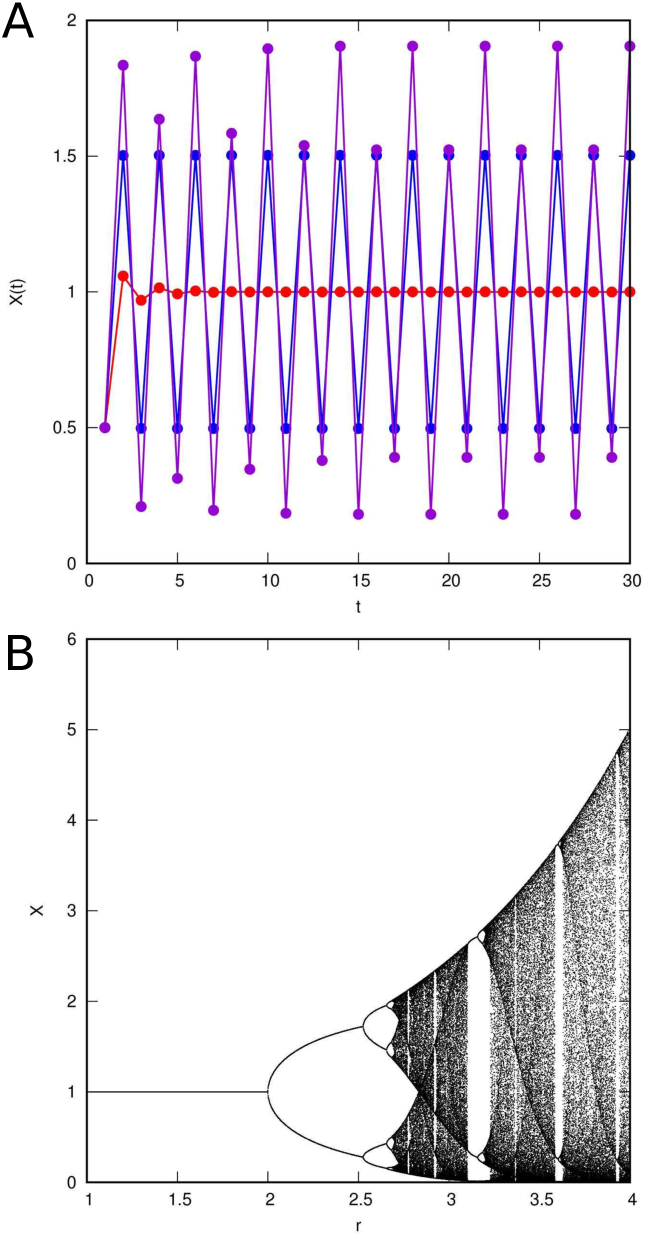
Single species dynamics in the model outlined in Equation 3. A - population trajectories when r equals 1.5 (red), 2.2 (blue) and 2.6 (purple). B - bifurcation diagram.

**Fig. 2.**
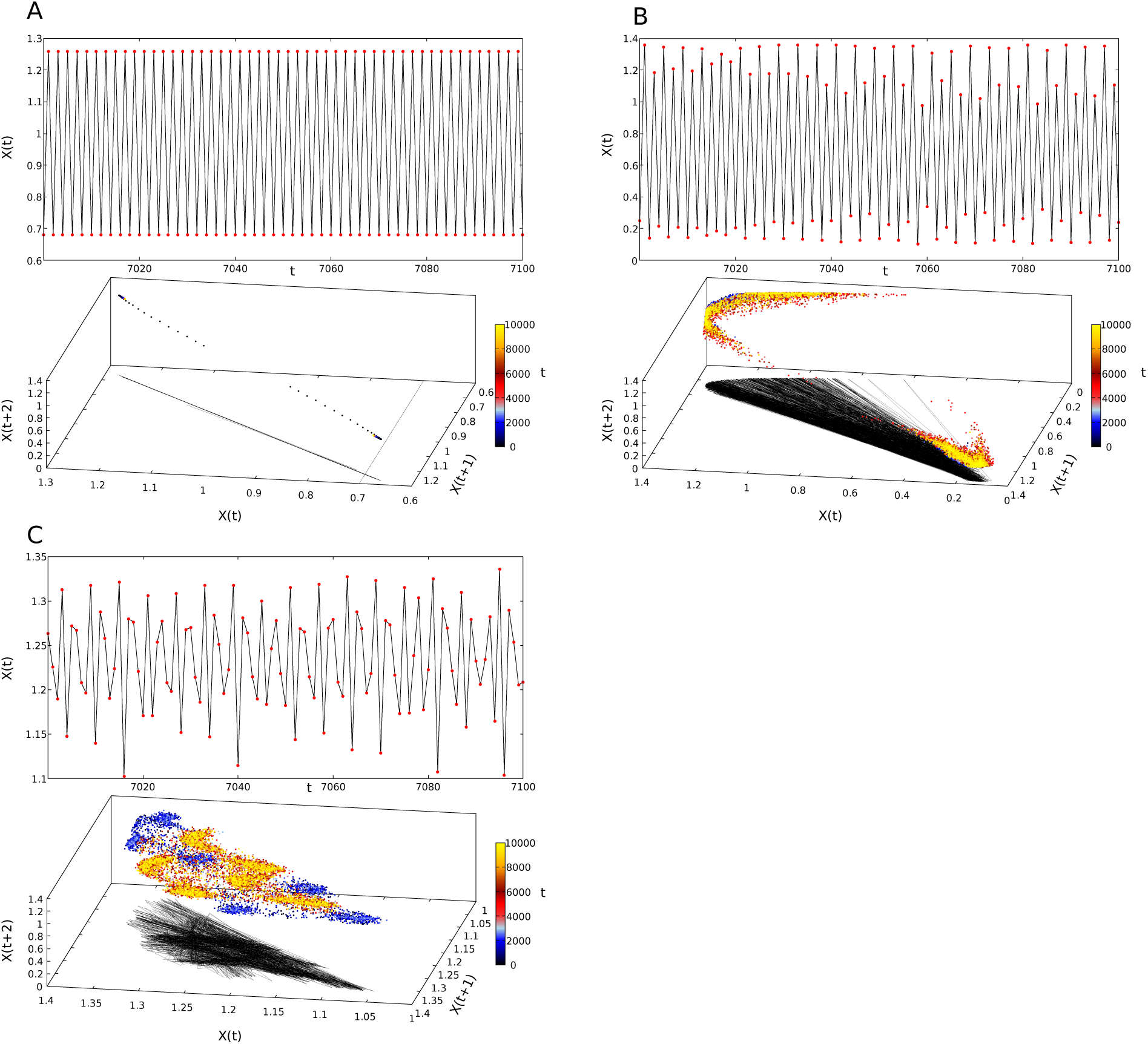
Population trajectories and phase plots (attractors) of taxa from the lower *Daptocephalus* Assemblage Zone (lDAZ), illustrating three types of dynamics. A - stable oscillation or limit cycle of a freshwater bivalve. B - quasi-periodic oscillation of a freshwater fish. C - chaotic dynamics of an omnivorous insect. Color bar illustrates time step (t). Upper plots show a subset of 10,000 time steps. Lines projected at base of attractor plots show the trajectories for the subsets.

**Fig. 3.**
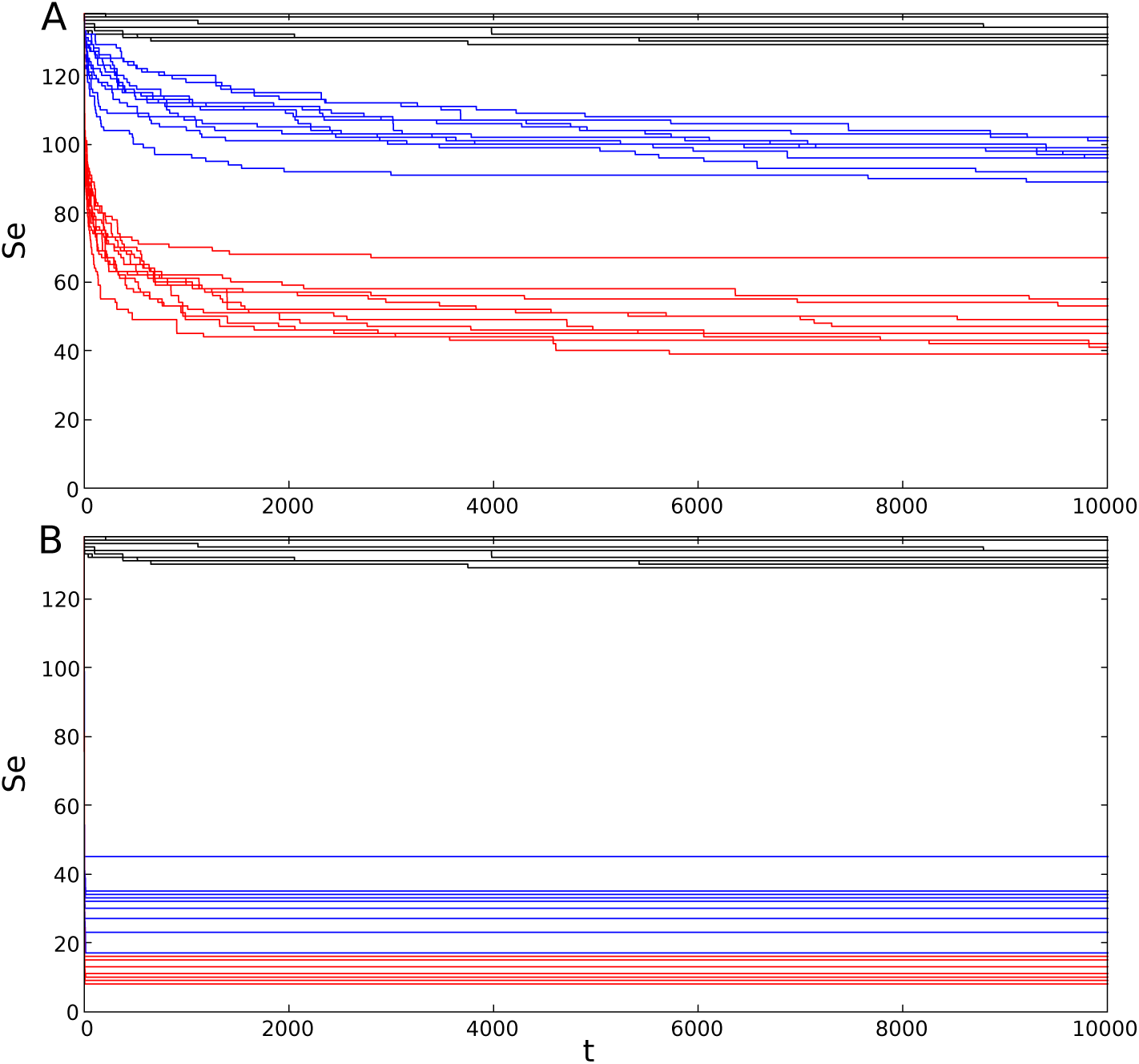
S declines as the model is iterated, asymptotically yielding Se. Se declines with increasing r and average negative interaction strength, as determined by the scaling factor c1. Both plots are of the lDAZ community, simulated at three different values of r and c1, with 10 food web simulations at each value.A - r equals 2 (black), 2.5 (blue) and 3 (red). c1=0.1. B - c1 equals 0.1 (black), 0.5 (blue), 1.0 (red). r=2.

**Fig. 4.**
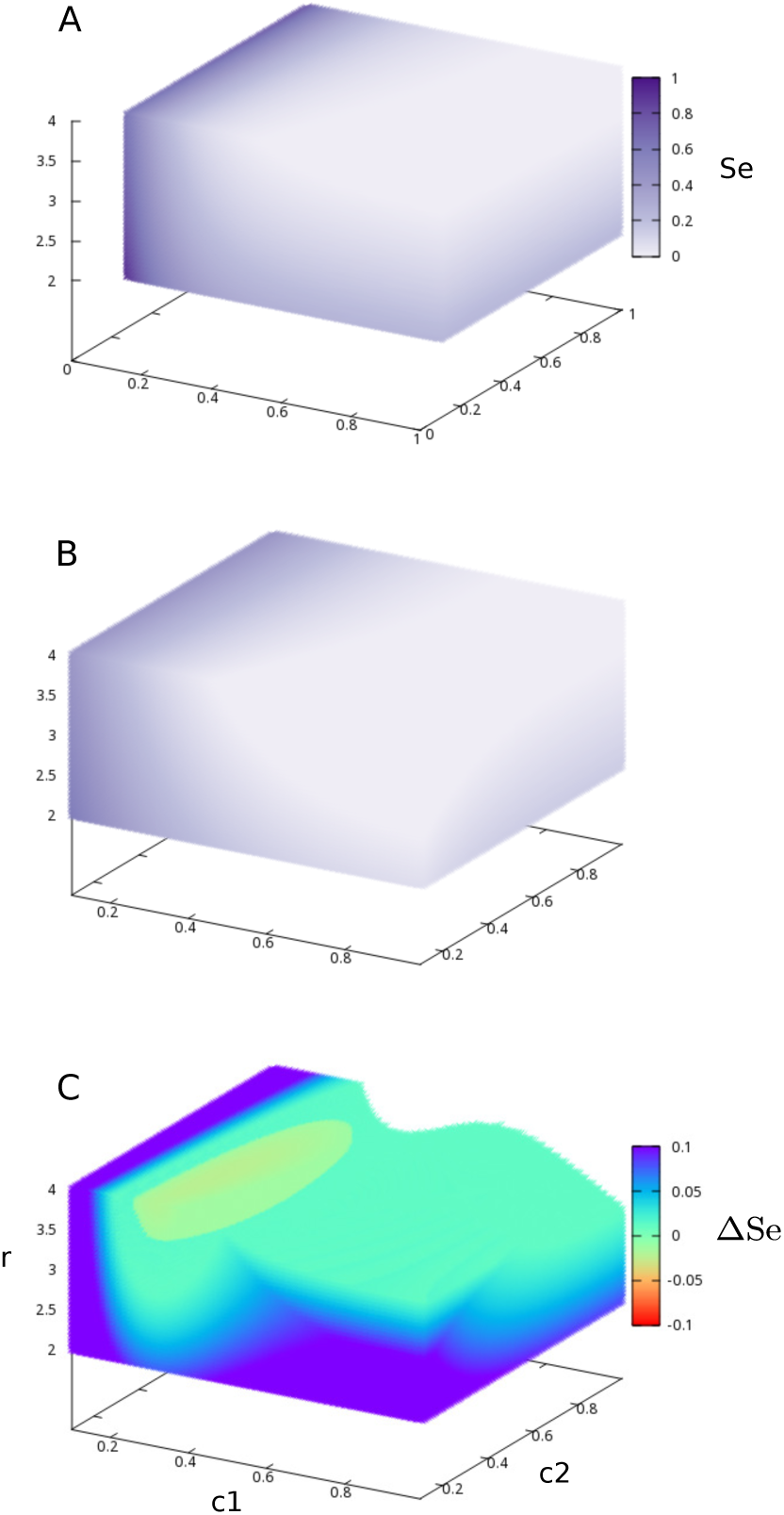
Visualization of Se and ΔSe throughout the r,c1,c2 parameter space, for the lDAZ community. Color intensity shows the value of Se in A and B. A - observed lDAZ community. B - random food web with S and the total number of predator-prey interactions equal to lDAZ’s S (S=138). C - ΔSe of the observed model minus the random model. Values of ΔSe=0 are not colored. Note that ΔSe ≥ 0 everywhere, except for a small region where r ≥ 3.5 and 0.2 ≤c1≤0.3.

**Table S1.** Guilds and guild richnesses of all paleocommunities. S is total richness.

| Guilds | IDAZ | uDAZ | Ph1 | Ph2 | Ph3 | LAZ | CAZ |
| --- | --- | --- | --- | --- | --- | --- | --- |
| Plants accessible to amniotes and arthropods | 1 | 1 | 1 | 1 | 1 | 1 | 1 |
| Aquatic microphytes | 1 | 1 | 1 | 1 | 1 | 1 | 1 |
| Aquatic macrophytes | 1 | 1 | 1 | 1 | 1 | 1 | 1 |
| Terrestrial plants accessible to arthropods only | 1 | 1 | 1 | 1 | 1 | 1 | 1 |
| Fish | 8 | 8 | 8 | 8 | 4 | 4 | 13 |
| Very small amphibians | 0 | 0 | 0 | 0 | 1 | 3 | 0 |
| Small amphibians | 0 | 0 | 0 | 0 | 0 | 3 | 0 |
| Medium amphibians | 1 | 1 | 0 | 0 | 0 | 0 | 1 |
| Large amphibians | 0 | 0 | 0 | 0 | 0 | 1 | 1 |
| Very large amphibians | 2 | 2 | 1 | 0 | 0 | 3 | 2 |
| Very small amniote herbivores | 4 | 4 | 0 | 0 | 3 | 4 | 10 |
| Very small amniote faunivores | 11 | 8 | 1 | 1 | 0 | 12 | 4 |
| Small amniote herbivores | 5 | 2 | 0 | 0 | 2 | 1 | 2 |
| Small amniote faunivores | 12 | 9 | 2 | 1 | 2 | 4 | 2 |
| Medium herbivores | 4 | 6 | 5 | 4 | 2 | 1 | 1 |
| Medium amniote carnivores | 9 | 5 | 1 | 1 | 4 | 2 | 1 |
| Large amniote herbivores | 5 | 4 | 2 | 1 | 0 | 0 | 0 |
| Large amniote carnivores | 3 | 5 | 0 | 0 | 0 | 0 | 0 |
| Very large amniote herbivores | 3 | 3 | 3 | 0 | 0 | 0 | 1 |
| Very large amniote carnivores | 2 | 3 | 1 | 0 | 0 | 0 | 1 |
| Molluscs | 2 | 2 | 1 | 0 | 0 | 1 | 0 |
| Conchostracans | 2 | 2 | 1 | 0 | 0 | 1 | 0 |
| Myriapods | 0 | 0 | 0 | 0 | 0 | 2 | 0 |
| Carnivorous insects | 8 | 8 | 2 | 1 | 7 | 8 | 7 |
| Herbivorous insects | 23 | 22 | 7 | 3 | 19 | 23 | 21 |
| Omnivorous insects | 30 | 29 | 20 | 9 | 24 | 29 | 28 |
| S | 138 | 127 | 59 | 33 | 72 | 106 | 99 |

**Table S2.**
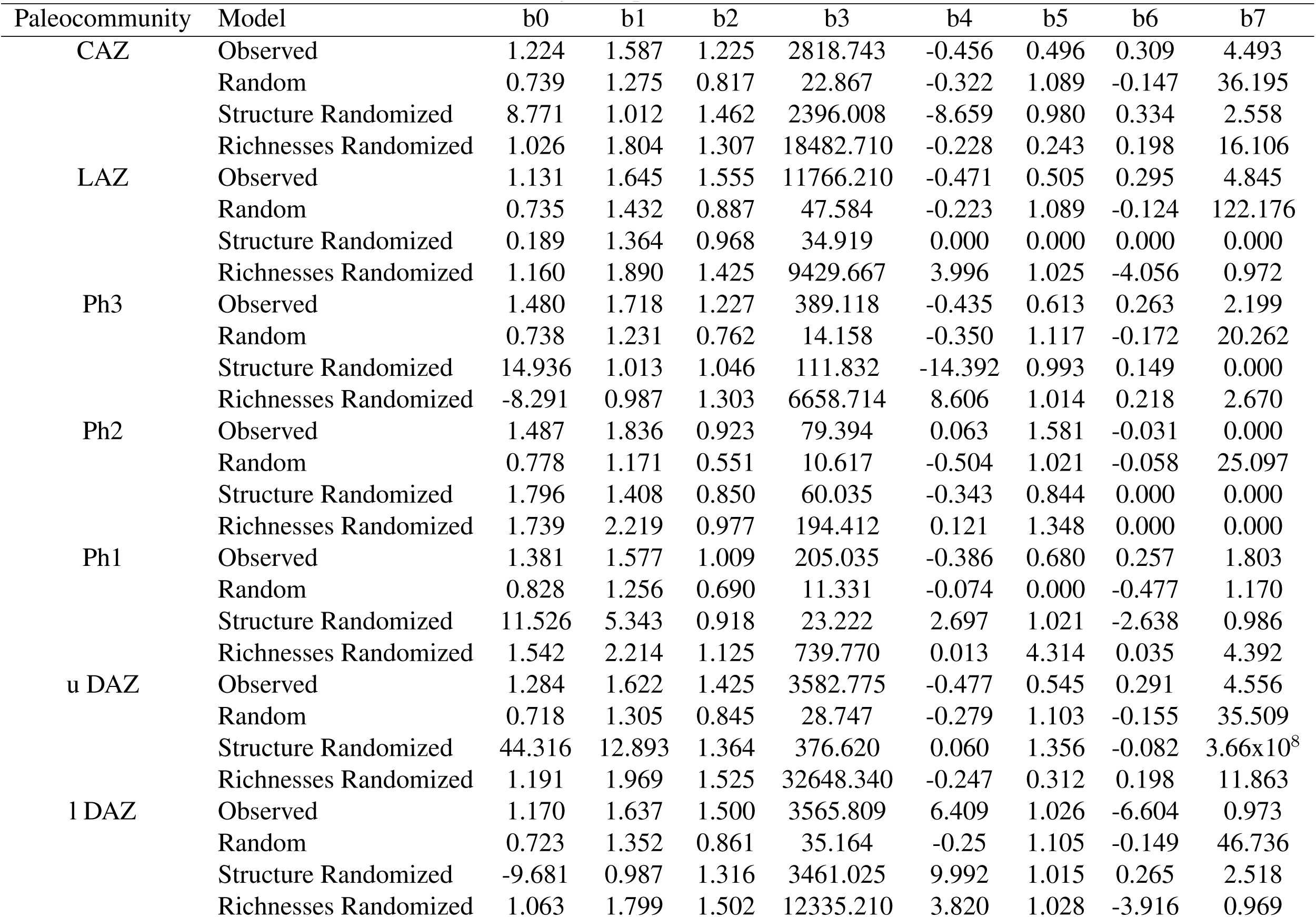
Coefficients for each model and community, in Equation 6.

| Paleocommunity | Model | b0 | b1 | b2 | b3 | b4 | b5 | b6 | b7 |
| --- | --- | --- | --- | --- | --- | --- | --- | --- | --- |
| CAZ | Observed | 1.224 | 1.587 | 1.225 | 2818.743 | -0.456 | 0.496 | 0.309 | 4.493 |
|  | Random | 0.739 | 1.275 | 0.817 | 22.867 | -0.322 | 1.089 | -0.147 | 36.195 |
|  | Structure Randomized | 8.771 | 1.012 | 1.462 | 2396.008 | -8.659 | 0.980 | 0.334 | 2.558 |
|  | Richnesses Randomized | 1.026 | 1.804 | 1.307 | 18482.710 | -0.228 | 0.243 | 0.198 | 16.106 |
| LAZ | Observed | 1.131 | 1.645 | 1.555 | 11766.210 | -0.471 | 0.505 | 0.295 | 4.845 |
|  | Random | 0.735 | 1.432 | 0.887 | 47.584 | -0.223 | 1.089 | -0.124 | 122.176 |
|  | Structure Randomized | 0.189 | 1.364 | 0.968 | 34.919 | 0.000 | 0.000 | 0.000 | 0.000 |
|  | Richnesses Randomized | 1.160 | 1.890 | 1.425 | 9429.667 | 3.996 | 1.025 | -4.056 | 0.972 |
| Ph3 | Observed | 1.480 | 1.718 | 1.227 | 389.118 | -0.435 | 0.613 | 0.263 | 2.199 |
|  | Random | 0.738 | 1.231 | 0.762 | 14.158 | -0.350 | 1.117 | -0.172 | 20.262 |
|  | Structure Randomized | 14.936 | 1.013 | 1.046 | 111.832 | -14.392 | 0.993 | 0.149 | 0.000 |
|  | Richnesses Randomized | -8.291 | 0.987 | 1.303 | 6658.714 | 8.606 | 1.014 | 0.218 | 2.670 |
| Ph2 | Observed | 1.487 | 1.836 | 0.923 | 79.394 | 0.063 | 1.581 | -0.031 | 0.000 |
|  | Random | 0.778 | 1.171 | 0.551 | 10.617 | -0.504 | 1.021 | -0.058 | 25.097 |
|  | Structure Randomized | 1.796 | 1.408 | 0.850 | 60.035 | -0.343 | 0.844 | 0.000 | 0.000 |
|  | Richnesses Randomized | 1.739 | 2.219 | 0.977 | 194.412 | 0.121 | 1.348 | 0.000 | 0.000 |
| Ph1 | Observed | 1.381 | 1.577 | 1.009 | 205.035 | -0.386 | 0.680 | 0.257 | 1.803 |
|  | Random | 0.828 | 1.256 | 0.690 | 11.331 | -0.074 | 0.000 | -0.477 | 1.170 |
|  | Structure Randomized | 11.526 | 5.343 | 0.918 | 23.222 | 2.697 | 1.021 | -2.638 | 0.986 |
|  | Richnesses Randomized | 1.542 | 2.214 | 1.125 | 739.770 | 0.013 | 4.314 | 0.035 | 4.392 |
| u DAZ | Observed | 1.284 | 1.622 | 1.425 | 3582.775 | -0.477 | 0.545 | 0.291 | 4.556 |
|  | Random | 0.718 | 1.305 | 0.845 | 28.747 | -0.279 | 1.103 | -0.155 | 35.509 |
|  | Structure Randomized | 44.316 | 12.893 | 1.364 | 376.620 | 0.060 | 1.356 | -0.082 | 3.66x10<sup>8</sup> |
|  | Richnesses Randomized | 1.191 | 1.969 | 1.525 | 32648.340 | -0.247 | 0.312 | 0.198 | 11.863 |
| l DAZ | Observed | 1.170 | 1.637 | 1.500 | 3565.809 | 6.409 | 1.026 | -6.604 | 0.973 |
|  | Random | 0.723 | 1.352 | 0.861 | 35.164 | -0.25 | 1.105 | -0.149 | 46.736 |
|  | Structure Randomized | -9.681 | 0.987 | 1.316 | 3461.025 | 9.992 | 1.015 | 0.265 | 2.518 |
|  | Richnesses Randomized | 1.063 | 1.799 | 1.502 | 12335.210 | 3.820 | 1.028 | -3.916 | 0.969 |

